# Landscape and evolutionary dynamics of terminal-repeat retrotransposons in miniature (TRIMs) in 48 whole plant genomes

**DOI:** 10.1101/010850

**Authors:** Dongying Gao, Yupeng Li, Brian Abernathy, Scott A. Jackson

**Affiliations:** Center for Applied Genetic Technologies, University of Georgia, 111 Riverbend Rd., Athens, GA 30602. USA

**Keywords:** plant, TRIM, retrotransposition, gene evolution, genomics

## Abstract

Terminal-repeat retrotransposons in miniature (TRIMs) are structurally similar to long terminal repeat (LTR) retrotransposons except that they are extremely small and difficult to identify. Thus far, only a few TRIMs have been characterized in the euphyllophytes and the evolutionary and biological impacts and transposition mechanism of TRIMs are poorly understood. In this study, we combined *de novo* and homology-based methods to annotate TRIMs in 48 plant genome sequences, spanning land plants to algae. We found 156 TRIM families, 146 previously undescribed. Notably, we identified the first TRIMs in a lycophyte and non-vascular plants. The majority of the TRIM families were highly conserved and shared within and between plant families. Even though TRIMs contribute only a small fraction of any plant genome, they are enriched in or near genes and may play important roles in gene evolution. TRIMs were frequently organized into tandem arrays we called TA-TRIMs, another unique feature distinguishing them from LTR retrotransposons. Importantly, we identified putative autonomous retrotransposons that may mobilize specific TRIM elements and detected very recent transpositions of a TRIM in *O. sativa.* Overall, this comprehensive analysis of TRIMs across the entire plant kingdom provides insight into the evolution and conservation of TRIMs and the functional roles they may play in gene evolution.

## Introduction

Retrotransposons are ubiquitous component of most eukaryotic genomes. These elements use an element-encoded mRNA as the transposition intermediate and can rapidly proliferate in copy number resulting in large differences in genome sizes between related species (Kumar and Bennetzen 1999; Piegu et al. 2006). Retroelement-induced mutations are usually stable and are widely used as molecular tools for gene-tagging and functional analysis (Hirochika 2001; Mazier et al. 2006). Retrotransposons provide raw material for evolutionary innovation, including new genes and gene regulatory networks and serve as essential components for centromere and telomere integrity (Chuong et al. 2013). Furthermore, retroelements can form functional genomic elements that regulate gene expression, maintain chromatin structure and contribute to histone modification and DNA methylation (Grewal and Jia 2007; ENCODE Project Consortium 2012).

LTR retrotransposons are the most plentiful mobile elements in plant kingdom. For example, there are more than 1.1 million LTR retroelements in maize, accounting for 75% of the genome (Schnable et al. 2009). LTR retrotransposons in plants are often large, up to 20 kb, and can have LTRs more than 5 kb in length (Kumar and Bennetzen 1999). These elements are often clustered into blocks greater than 100-kb via multiple-layers of nested insertions (Kronmiller and Wise 2008). Moreover, LTR retrotransposons can have distinct chromosomal distribution patterns. For example, LTR retrotransposons can be found in intergenic regions but are most concentrated in highly heterochromatic regions (Jiang et al. 2002; Jiang et al. 2003; Ammiraju et al. 2007). Plant LTR retrotransposons are very dynamic and except for a few examples, e.g. centromeric retrotransposons (CRs) in grasses (Lee et al. 2005; Piegu et al. 2006; Gao et al. 2009), it is nearly impossible to detect sequence homology between related species.

Terminal repeat retrotransposons in miniature (TRIMs) are unusual elements that maintain similarities with LTR retrotransposons, including terminal direct repeats and target site duplication (TSD), but are small, less than 1000 bp (Witte et al. 2001) and as small as 292 bp (Gao et al. 2012), and do not encode proteins needed for movement. Owing to their extremely short length and lack of capacity to encode proteins, TRIMs are difficult to annotate. To date, only ten TRIM families, *Katydid*-At1, At2, At3 (Witte et al., 2001), Br1-Br4, *Katydid*-At4 (Yang et al., 2007), Cassandra (Kalendar et al. 2008; Yin et al. 2014) and SMART (Gao et al. 2012), have been reported in the euphyllophytes. Recently, a TRIM was reported in the red harvester ant (PbTRIM, *Pogonomyrmex barbatus*, Zhou and Cahan 2012), the only one reported in animals and microbes. Most of these studies have focused on one or a few TRIM families and no TRIM elements have been found in lycophytes and non-vascular plants. Thus, the evolutionary impacts of TRIMs on host genomes and mechanisms involved in their emergence and disappearance remain poorly understood. Due to the availability of more plant genome sequences, we are now able to analyze and compare TRIMs across a broad evolutionary range of species.

To identify new TRIMs and understand their evolution and mechanism of transposition, we analyzed 48 whole genome sequences, including spermatophytes (seed plants), lycophyte, bryophyta and algae. We identified complete TRIM elements in all the flowering plants and, for first time, in a lycophyte and non-vascular plants. Based on comparative analyses, we grouped TRIMs into 156 families of which 146 are new TRIM families. We determined the insertion times and organization of TRIMs and found that they are frequently tandemly arranged. We observed that TRIMs are enriched in genic regions and likely play a role in gene evolution. Importantly, we identified the first putative autonomous LTR retrotransposons for a TRIM and uncovered recent transpositions of a TRIM family in *O. sativa*. These results provide a better understanding of the evolutionary dynamics of plant genomes and the role that TRIM elements play in plant genome and gene evolution.

## Results

### Identification and abundance of plant TRIMs

Combining *de novo* and homology-based searches, we analyzed 48 plant genomes available as of April 1, 2013 (Supplemental Table S1). We identified TRIM transposons in 43 genomes, including all 30 eudicots and nine monocots. Notably, we found TRIMs in the lycophyte, *S. moellendorffii*, and three algae genomes, *C. reinhardtii*, *V. carteri* and *C. crispus* (Table 1). To our knowledge, this is the first time that TRIMs have been reported in lycophytes and non-vascular plants. However, TRIM elements were not found in *P. patens*, and four other algae genomes, *C. variabilis, O. lucimarinus, O. tauri* and *C. merolae*.

**Table 1.** Summary of TRIMs in 43 sequenced plant genomes

| Plant genome | Genus/Family of plant | Number of TRIM subfamily |  |  |  | Copy number |  | Fraction (%) |
| --- | --- | --- | --- | --- | --- | --- | --- | --- |
|  |  | Shared between families | Family specific | Species specific | Total | Complete | Total |  |
| Tomato (*Solanum lycopersicum*) | *Solanum/Solanaceae* | 6 | 4 |  | 10 | 560 | 9,162 | 0.32 |
| Currant Tomato (*Solanum pimpinellifolium*) | *Solanum/Solanaceae* | 7 | 4 |  | 11 | 178 | 10,199 | 0.29 |
| Potato (*Solanum tuberosum*) | *Solanum/Solanaceae* | 4 | 5 |  | 9 | 451 | 12,473 | 0.46 |
| Cucumber (*Cucumis sativus*) | *Cucumis/Cucurbitaceae* | 4 |  |  | 4 | 30 | 2,816 | 0.21 |
| Maskmelon (*Cucumis melo*) | *Cucumis/Cucurbitaceae* | 3 |  |  | 3 | 44 | 2,072 | 0.09 |
| Watermelon (*Citrullus lanatus*) | *Citrullus/Cucurbitaceae* | 5 |  |  | 5 | 228 | 4,779 | 0.21 |
| Plum blossom (*Prunus mume*) | *Prunus mume/Rosaceae* | 5 | 2 |  | 7 | 83 | 5,719 | 0.47 |
| Apple (*Malus x domestica*) | *Malus/Rosaceae* | 7 |  |  | 7 | 2,043 | 25,835 | 0.74 |
| Pear (*Pyrus bretschneideri*) | *Pyrus/Rosaceae* | 6 | 1 |  | 7 | 2,286 | 20,092 | 1.26 |
| Strawberry (*Fragaria vesca*) | *Fragaria/Rosaceae* | 4 |  |  | 4 | 132 | 1,605 | 0.18 |
| Marijuana (*Cannabis sativa*) | *Cannabis/Cannabaceae* | 5 |  | 1 | 6 | 362 | 14,147 | 0.55 |
| Lotus (*Lotus japonicus*) | *Lotus/Fabaceae* | 6 | 2 | 1 | 9 | 379 | 7,943 | 0.88 |
| Barrel medic (*Medicago truncatula*) | *Medicago/Fabaceae* | 7 | 1 |  | 8 | 46 | 8,416 | 0.56 |
| Chickpea (*Cicer arietinum*) | *Cicer/Faboideae* |  | 2 |  | 2 | 102 | 1,499 | 0.21 |
| Soybean (*Glycine max*) | *Glycine/Faboideae* | 9 | 1 | 6 | 16 | 261 | 10,102 | 0.25 |
| Pigeonpea (*Cajanus cajan*) | *Cajanus/Faboideae* | 9 | 3 | 1 | 13 | 840 | 20,915 | 0.67 |
| Sanskrit (*Jatropha curcas*) | *Jatropha/Euphorbiaceae* | 5 |  | 2 | 7 | 177 | 3,390 | 0.28 |
| Flax (*Linum usitatissimum*) | *Linum/Linaceae* | 3 |  | 2 | 5 | 71 | 4,149 | 0.33 |
| Castor bean plant (*Ricinus communis*) | *Ricinus/Euphorbiaceae* | 2 |  |  | 2 | 90 | 385 | 0.02 |
| Poplar (*Populus trichocarpa*) | *Populus/Salicaceae* | 5 |  | 1 | 6 | 839 | 5,292 | 0.28 |
| Thale cress (*Arabidopsis thaliana*) | *Arabidopsis/Brassicaceae* | 5 | 3 |  | 8 | 36 | 876 | 0.09 |
| Lyrate rockcress (*Arabidopsis lyrata*) | *Arabidopsis/Brassicaceae* | 9 | 5 |  | 14 | 259 | 1,724 | 0.25 |
| Pallus (*Thellungiella salsuginea*) | *Thellungiella/Brassicaceae* | 9 | 1 |  | 10 | 98 | 1,406 | 0.16 |
| Turnip mustard (*Brassica rapa*) | *Brassica/Brassicaceae* | 9 | 1 |  | 10 | 269 | 3,030 | 0.26 |
| Eutrema parvulum (*Thellungiella parvula*) | *Eutrema/Brassicaceae* | 3 | 1 |  | 4 | 25 | 539 | 0.10 |
| Papaya (*Carica papaya*) | *Carica/Caricaceae* |  |  | 1 | 1 | 5 | 897 | 0.09 |
| Cocoa (*Theobroma cacao*) | *Theobroma/Malvaceae* |  |  | 1 | 1 | 45 | 360 | 0.03 |
| Cotton (*Gossypium raimondii*) | *Gossypium/Malvaceae* | 1 | 2 |  | 3 | 19 | 19,008 | 0.35 |
| Grape (*Vitis vinifera*) | *Vitis/Vitaceae* | 4 |  | 1 | 5 | 228 | 8,890 | 0.34 |
| Sweet Orange (*Citrus sinensis*) | *Citrus/Rutaceae* | 2 |  | 1 | 3 | 30 | 1,180 | 0.09 |
| Sorghum (*Sorghum bicolor*) | *Sorghum/Poaceae* | 1 | 5 |  | 6 | 282 | 2,922 | 0.11 |
| Maize (*Zea mays*) | *Zea/Poaceae* | 1 | 3 | 3 | 7 | 1,361 | 9,036 | 0.12 |
| Foxtail (*Setaria italica*) | *Setaria/Poaceae* | 1 | 4 |  | 5 | 129 | 1,032 | 0.07 |
| Rice-Japonica (*Oryza sativa, Japonica*) | *Oryza/Poaceae* | 2 | 9 |  | 11 | 379 | 2,911 | 0.18 |
| Rice-*Indica* (*Oryza sativa*, *Indica*) | *Oryza/Poaceae* | 2 | 9 |  | 11 | 364 | 3,252 | 0.19 |
| Brachyantha (*Oryza brachyantha*) | *Oryza/Poaceae* | 2 | 8 | 1 | 11 | 116 | 1,506 | 0.15 |
| Purple false brome (*Brachypodium distachyon*) | *Brachypodium/Poaceae* | 1 | 2 | 2 | 5 | 75 | 1,685 | 0.15 |
| Date palm (*Phoenix dactylifera*) | *Phoenix/Arecaceae* | 2 |  | 7 | 9 | 777 | 13,358 | 0.78 |
| Banana (*Musa acuminata*) | *Musa/Musaceae* |  |  | 2 | 2 | 126 | 4,577 | 0.23 |
| Spikemoss (*Selaginella moellendorffii*) | *Selaginella/Selaginellaceae* | 2 |  | 6 | 8 | 1,177 | 10,158 | 1.19 |
| Green alga (*Chlamydomonas reinhardtii*) | *Chlamydomonas/Chlamydomonadaceae* | 1 |  | 5 | 6 | 31 | 1,349 | 0.21 |
| Volvox (*Volvox carteri*) | *Volvox/Volvocaceae* |  |  | 5 | 5 | 292 | 2,052 | 0.27 |
| Irish moss (*Chondrus crispus*) | *Chondrus/Gigartinaceae* |  |  | 3 | 3 | 75 | 422 | 0.09 |
| **Total** |  | 159 | 78 | 52 | 289 |  |  |  |

The conservation of TRIM elements across species was reported previously (Witte et al. 2001; Kalendar et al. 2008; Gao et al. 2012). Thus, TRIM elements from all 43 genomes were grouped into TRIM subfamilies rather than families. A total of 289 TRIMs subfamilies were identified with an average of 6.7 subfamilies per genome (Table 1), varying from 16 in *G. max* to one in *C. papaya* and *T. cacao*.

Even though TRIMs are small, the terminal direct repeats are still referred to as long terminal repeats (LTRs) for comparison to other LTR retrotransposons. The average size of the 289 subfamilies was 685 bp, much smaller than typical plant LTR retroelements (4-10 kb on average, Wessler 2006). The average LTR of a TRIM was 198 bp. Among the 289 subfamilies, 225 (77.9%) were smaller than 1000 bp and the LTR sizes of 197 (68.1%) were less than 250 bp (Supplemental Figure S1A, B). The size of a TRIM, CcaRetroS9, in *C. cajan* was only 233 bp with 52-bp LTRs and is the smallest TRIM identified thus far.

The copy numbers of TRIMs were highly variable between genomes. The majority (65%, 28/43) of the plants harbored more than 2,000 complete or fragmented TRIMs, only six (14%) had fewer than 1,000 TRIMs (Table 1). Most, 174 of the 289 subfamilies (60%), had copy numbers less than 500, and about one-quarter (70/289) had copy numbers greater than 1,000 (Supplemental Figure S1C). Some subfamilies were highly abundant, for example GraRetroS4 was present in ˜18,000 copies in *G. raimondii*.

### Conservation and comparison of TRIMs

To determine the phylogenetic distribution and group the TRIM elements, all 289 TRIM subfamilies were used to search GenBank and conduct all-by-all BLASTN searches. 159 subfamilies were found in more than two plant taxonomic families, 78 subfamilies were present in multiple genomes from a same plant family, termed ‘family-specific TRIMs’, whereas 52 subfamilies were found only in a single genome, termed ‘species-specific TRIMs’. Species-specific TRIMs may have homologs that were either lost or diverged in other genomes or not represented in GenBank (Table 1).

The TRIMs from the 43 plants were then grouped into families based on sequence similarity. A total of 156 TRIM families were identified, 60 of which were shared between plant families, 44 were specific to single plant family, and 52 were species-specific. Of note, 146 TRIM families were identified for the first time. We also found new members for the 10 previously reported TRIM families (Witte et al., 2001; Yang et al., 2007; Kalendar et al. 2008; Gao et al. 2012), such as complete Cassandra transposons in *C. sativa* and other plants.

The TRIMs from three plant taxonomic families, the Legumes (*Fabaceae*), Cruciferae (*Brassicaceae*) and Grasses (*Poaceae*), are detailed in Figure 1. These three families were chosen as each contains more than five sequenced genomes, contain both dicots and monocots and represent ˜140-150 million years (My) of evolution (Chaw et al. 2004). They provide a resource to analyze the conservation and speciation of plant TRIMs. Additionally, the grasses and legumes contain many crops and are the two most economically and agriculturally important plant families.

**Figure 1.**
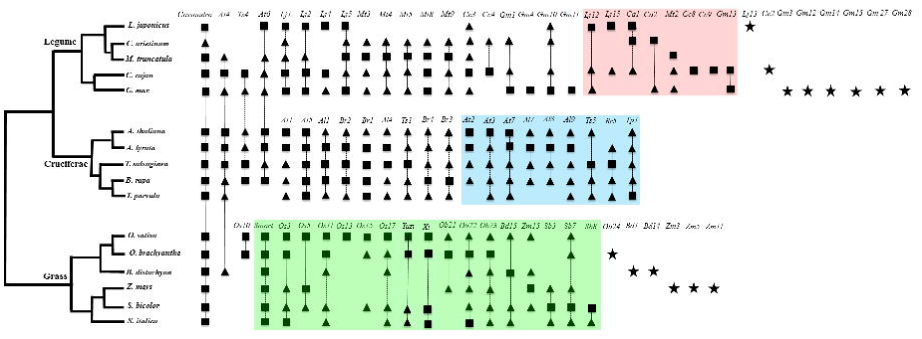
Comparison of TRIMs in three plant taxonomic families. Black squares and triangles represent complete and fragmented TRIMs, respectively, shared within and between plant genomes. Black stars indicate TRIMs present in a single genome. TRIMs grouped into a single family are linked by dashed lines. TRIMs in the pink, blue and green boxes are present only in legumes, cruciferae and grasses, respectively.

Within the Cruciferae *A. lyrata* and *B. rap*a shared a common ancestor with the model plant *A. thaliana* about 13 and 43 million year ago (Mya), respectively (Beilstein et al. 2010). Nine TRIM families were previously reported in this plant family, including At1-4 and Cassandra in *A. thaliana* (Witte et al. 2001; Yang et al. 2007; Kalendar et al. 2008) and Br1-4 in *B. rap*a (Yang et al. 2007). We identified 13 new TRIM families. Among the 22 TRIM families, two, Cassandra and At4, have complete or fragmented homologs in legumes and grasses, 11 were shared between the Cruciferae and other dicots, and nine families were found only within the Cruciferae (Figure 1).

36 TRIM families were found in the five legume genomes including two published elements, Cassandra and At4. Among the 36, 15 were shared between legumes and other plant families, two families, GmaRetroS4 (abbreviated as *Gm4*) and GmaRetroS11 (*Gm11*) from *G. max*, were absent in the other four legumes but homologs were found in other plants. For instance, a homolog of *Gm4* was found in *Trifolium pratense* (GeneBank accession number: BB931325:395-501, E value=2 X e^-7^). Eight family-specific TRIMs, LjaRetroS12, 15, CarRetroS1, 2, MtrRetroS2, CcaRetroS8, 9 and GmaRetroS13, were found in subsets of the five sequenced legumes that last shared a common ancestor about 50 Mya (Lavin et al. 2005). Eight species-specific TRIM families were found within a single legume genome.

In addition to the two previously described TRIM families, Cassandra (Kalendar et al. 2008) and SMART (Gao et al. 2012), 23 new families were identified within the grasses. Family OsaRetroS10 (*Os10*) had complete elements in *O. sativa* and *O. brachyantha* and homologs were found in *S. lycopersicum* (AC243477:1845-1967, E value=7 X e^-^8^^) and *S. pimpinellifolium* (AGFK01075962: 4312-4434, E value=7e^-11^). Ten TRIM families identified in *O. sativa* and *O. brachyantha* have complete and/or fragmented copies in *Z. mays* and/or *S. bicolor* that diverged from the *Oryza* genus ˜50-80 Mya (Gaut 2002). One, two and three TRIM families were found only in *O. brachyantha*, *B. distachyon* and *Z. mays*, respectively.

### TRIM-mediated gene evolution

TRIMs have been postulated to be involved in gene divergence and regulation (Witte et al. 2001; Kalendar et al. 2008; Gao et al. 2012). However, these focused on only one or a few TRIM families and did not provide a global, genome-wide view of the impact of TRIMs on gene evolution and function. Therefore, we examined the distribution of TRIMs with respect to genes in 14 of the plant genomes. Our data indicate that TRIMs are enriched in genic regions as 18.8-49.4% of TRIMs are located in or near (1.5 Kb upstream) genes (Supplemental Table S2). Interestingly, an average of 2.7% of the TRIMs within a genome have been recruited as exons, including coding DNA sequences (CDS) and untranslated regions (UTRs). A previous study revealed that ∽45% of the TRIMs were present within or near predicted genes in red harvester ant (Zhou and Cahan 2012). All these data indicate that TRIMs may exhibit similar insertion preferences (higher frequency in/near genes), in both plants and animal, and suggests that TRIMs may play an important role in the gene diversification.

To provide more insight into the contribution of TRIMS to gene function and divergence, we carefully analyzed two genomes, *G. max* and *Z. mays*, and compared gene structures of TRIM-related genes, those that contain TRIM sequences, and non-TRIM-related genes. In both genomes, TRIM-related genes had more exons and were larger than non-TRIM-related genes (Figure S2; Supplemental Table S3). For example, in *G. max* the average exon number of TRIM-related genes was 12.2 versus 5.9 for non-TRIM-related genes. Differences in exon number, exon and intron sizes between TRIM-related and non-TRIM-related genes were statistically significant for both species, p-values from two sample T-tests after log transformation were less than 2.2 X 10^-16^.

Because TRIMs are small, average size (total coverage/total number of complete and fragmented TRIMs) of 181.5 bp and 271.9 bp in *G. max* and *Z. mays,* respectively, we expected relatively little contribution to the expansion of genes. Thus, the large differences in exon number and gene size may reflect an insertional bias of TRIMs into larger genes. To test this hypothesis, TRIM-related and non-TRIM-related genes in the two genomes were used to find orthologous genes in their closest relatives: *C. cajan* and *P. vulgaris* for *G. max*, diverged ∽20 and 15 Mya, respectively (Lavin et al. 2005), and *S. bicolor* and *O. sativa* for *Z. mays*, diverged ∽10 and 50-80 Mya, respectively (Gaut 2002). Results indicated that in all four genomes that homologs of TRIM-related genes also have higher exon numbers and are larger than orthologs of non-TRIM-related genes. The exon number and sizes of TRIM-related and non-TRIM-related genes were similar to their orthologous (Supplemental Table S4). However, the introns of both TRIM-related and non-TRIM-related genes in *Z. mays* were larger than orthologs from *S. bicolor* and *O. sativa*, likely due to higher transposon density in *Z. mays* (Schnable et al. 2009). We also observed that TRIM-related genes are more conserved than non-TRIM-related genes, 95.4% (2383/2497) of TRIM-related genes in *G. max* had homologs in both *C. cajan* and *P. vulgaris*, whereas, only 42.9% (22863/53290) of non-TRIM-related genes had homologs in both relatives. Taken together, our results indicate that TRIMs preferentially insert or are retained in large genes.

Transposons represent an important pathway for functional divergence of duplicated genes and emergence of new genes in both animals and plants (Long et al. 2013). Since TRIMs are enriched in genic regions and contribute to exons (Supplemental Table S2), we were interested to determine the effect of TRIM insertions on the divergence or evolution of host genes. To do this, TRIM-related genes were used to search GenBank and annotated genes from *G. max* and *Z. mays*. 15 TRIM-related genes (12 with expression support) in *G. max* have no homolog in any other genomes, and in *Z. mays* 35 TRIM-related genes (32 with expression support) were specific to *Z. mays*. By comparing genes with TRIM-derived exons to homologous genes from other species, we found that TRIMs likely resulted in creating new or alternatively spliced exons. For example, LOC100815590 from *G. max* exhibits ∽ 92% DNA sequence identity to its paralogous gene LOC100778729, however, LOC100815590 contains a solo-LTR from TRIM GmaRetroS12 recruited as the second exon and encodes 121-aa protein that is diverged and shorter than the predicted proteins from homologous genes from *G. max, M. truncatula* and *C. sinensis* (Supplemental Figure S3). The ratio of the nonsynonymous (Ka) and synonymous (Ks) substitutions rate between LOC100815590 and LOC100778729 was 1.5 indicating positive selection has acted affecting gene evolution. TRIM insertions may also affect the UTRs. We identified three instances, one in *Z. mays* and two in *G. max*, in which a complete TRIM element was located in 3’UTR region resulting in UTR sizes larger than that found in homologous genes (Supplemental Figure S4).

To further explore the functional consequences of TRIM insertions into genes, all annotated genes in *G. max* and *Z. mays* were used to query the protein sequences of *A. thaliana* from the TAIR10 genome release (https://www.arabidopsis.org). From this, 48,851 genes in *G. max* and 63,540 genes in *Z. mays* were assigned to Gene Ontologies (GO) and all these genes were classified into three categories including molecular function, cellular component and biological process. All GO categories contained TRIM-related genes (Supplemental Table S5). We did observe that TRIM-related genes were enriched for hydrolase activity and underrepresented in unknown molecular functions under the molecular function category, as compared to non-TRIM related genes for both *G. max* and *Z. mays*. In the biological process classification, TRIM-related genes encoding cell organization and biogenesis were more frequent than non-TRIM-related genes. These trends are different from that in *Mus musculus* where LTR retrotransposons are preferentially associated with genes encoding metabolic functions (DeBarry et al. 2006).

### Gene acquisitions related to TRIMs

Transposon-based gene capture is an important mechanism for gene evolution (Kapitonov and Jurka 2003; Jiang et al. 2004). Only one TRIM-mediated gene acquisition event has been reported to date, in *A. thaliana* (Witte et al. 2001). To assess the incidence of TRIM-based gene capture, the 289 TRIM subfamilies were used for BLASTN and BLASTX searches to detect the significant alignments (E value < 1 X 10^-10^) to expressed genes. From this, 30 TRIM elements from seven subfamilies contained gene fragments including one in *M. truncatula* and six in *G. max* (Supplemental Table S6). The sizes of the TRIMs range from 1172 to 1449 bp, similar to that of PACK-MULEs in rice (∽1.5 Kb, Jiang et al. 2004), and their internal regions had more than 70% sequence identity to the host genes. All these TRIMs contain only transcribed exon fragments, no introns. Two TRIMs carried exons from more than two genes. For instance, the internal region of GmaRetroS15 contains 217-bp and 160-bp sequences highly identical to the 5’UTR of LOC10081263 and an exon of LOC100820519, respectively. It also carries a 346-bp fragment with 76% sequence identity to the 5-9^th^ exons, but no introns, of LOC100798768, annotated as casein kinase I isoform delta-like protein (Fig. 2). These data suggest that the TRIM-mediated gene acquisition may differ from DNA transposons, such as PACK-MULEs that contain both exons and introns of cellular genes (Jiang et al. 2004; Juretic et al. 2005), and are more similar to an LTR retrotransposon, Bs1 in maize, which captured exons only (Bureau et al. 1994; Jin and Bennetzen 1994; Elrouby and Bureau 2010), and non-LTR retrotransposon L1 in human (Goodier et al. 2000).

**Figure 2.**
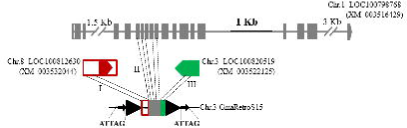
Gene acquisitions related to TRIM GmaRetroS15 in *G. max*. Black triangles and arrows denote TRIM LTRs and TSDs, respectively. Solid boxes and lines are exons and introns of three genes marked with different colors. The pentagons are the last exons of the genes and indicate transcription orientation. I, II and III indicate the fragments from three host genes. The cDNA sequence for each gene model is shown in parenthesis.

Among the 30 elements carrying gene fragments, all have two or more copies except GmaRetroS1 and GmaRetroS28, (Supplemental Table S6), all the elements contained both LTRs, and were flanked by 5-bp TSDs. One complete copy each was found for GmaRetroS1 and GmaRetroS28 in *G. max,* although other nearly complete copies were found. This suggests that following gene acquisition, additional transposition events occurred resulting in increased copy numbers.

### Tandemly arrayed (TA)-TRIMs

A typical LTR retrotransposon contains 5’ and 3’ LTRs flanking an internal region that often encodes proteins required for retrotransposition. We refer to this structure as L_2_I_1_ where L_2_ and I_1_ refer to two **<u>L</u>**TRs and an **<u>I</u>**nternal sequence, respectively. In addition to the typical TRIM elements (L_2_I_1_), some TRIMs arranged tandemly and contained more than three LTRs and two internal regions, hereafter, referred to as tandemly arrayed (TA)-TRIMs. So far, this peculiar structure was reported only for the Cassandra TRIM which LTRs contain sequences similar to cellular 5S rRNA that is also tandemly arranged (Kalendar et al. 2008; Yin et al. 2014). However, 5S rRNA sequences were not found in other any TRIM families.

To determine the incidence of TA-TRIMs, the 289 TRIM subfamilies were used to search the 43 genomes and the resulting sequences were manually inspected to determine their structures and TSDs. TA-TRIMS seem to be common in plant genomes as 129 subfamilies were found to have TA-TRIM sequences in 35 of the 43 genomes (Supplemental Table S7). There was variation among TRIM families for TA-TRIM formation. For example, complete Cassandra TRIMs were found in 17 plant species (15 shown in Figure 1 and *C. sativa* and *J. curcas*). Tandemly arrayed Cassandra elements were present in 15 of the 17 genomes, excepting *G. max* and *B. distachyon*. This contrasts to other families such as Xi in the grasses where no TA-TRIMS were found.

To gain more insight into TA-TRIMs within a single species, we focused on the maize genome there were 93 tandem arrays from four TRIM subfamilies. These arrays varied in organization and contained varying numbers of LTRs and internal sequences, such as three LTRs and two internal regions (L_3_I_2_), and five LTRs and four internal regions (L_5_I_4_) (Figure 3, Supplemental Table S8). Among all the TA-TRIMs identified in maize, L_3_I_2_ was the most frequent, accounting for more than 67% (63/93) of all TA-TRIMs. To validate TA-TRIMs in maize, we conducted PCR analysis using primers that targeted regions flanking TA-TRIMs from the Zma-SMART subfamily (Figure 3), and further confirmed by sequencing. This validated the structure and organization of the TA-TRIMS in that they were not artifacts of errors in genome assembly.

**Figure 3.**
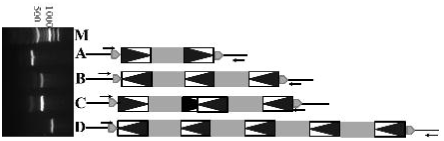
TA-TRIMs of Zma-SMART in maize genome. Boxes containing black triangles indicate the LTRs of TRIMs and grey boxes denote the internal regions of TRIMs. The grey pentagons are TSDs that flank TRIMs and arrows indicate the PCR primers used to validate the TRIM sequences. M means 100 bp DNA ladder; A indicates a typical Zma-SMART TRIM with two LTRs and one internal region (AC186328: 154584-154863; TSD: AACAT); B indicates a TA-TRIM with three LTRs and two internal regions (AC210283: 61391-61889; TSD: GGGTT); C indicates a TA-TRIM with two inverted TRIMs (AC220956: 117725-118283; TSD: CTTCA); and D indicates a TA-TRIM with five LTRs and four internal regions (AC185340: 80554-81415; TSD: ATAAT).

### Estimations of TRIM insertion times

To gain insight into the evolutionary dynamics of TRIM elements, we calculated the insertion times of TRIMs for six plant genomes, *A. thaliana*, *G. max*, *P. bretschneideri*, *O. sativa*, *S. moellendorffii* and *V. carteri*, which cover dicots, monocots, lycophyte, and algae. For comparison, the integration dates of both Ty1-copia and Ty3-gypsy retroelements were calculated (Figure 4). The Ty1-copia and Ty3-gypsy transposons exhibit similar amplification patterns in the six genomes with bursts of transposition less than 1 million year (Myr) and most, 80%, Ty1-copia and Ty3-gypsy elements less than 3 Myr. These results are consistent with the previous results from *A. thaliana* (Pereira 2004), *O. sativa* (Vitte et al. 2007) and *G. max* (Du et al. 2010). Furthermore, less than 2.9% of Ty1-copia and Ty3-gypsy elements had insertion times greater than 5 Myr, probably due to mutation and elimination (Vitte et al. 2007).

**Figure 4.**
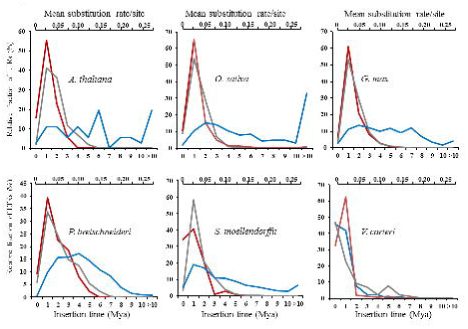
Estimation of insertion times of LTR retrotransposons and TRIMs from six plant species. Red, grey and blue lines represent Ty3-gypsy, Ty1-copia and TRIMs, respectively.

In contrast to Ty1-copia and Ty3-gypsy retrotransposons, TRIMs exhibited distinct amplification patterns. Only minor amplification peaks were seen in *A. thaliana*, *G. max*, *O. sativa*, and *S. moellendorffii*; whereas, a large amplification peak, ∽4 Mya, was observed in *P. bretschneideri*. Thus, it seems that amplification of TRIMs occurred over a longer time period than other TEs, especially as we found numerous TRIMs older than 10 Myr. Only 10-25% of TRIMs were younger than 1 Myr which indicates that TRIMs have likely been suppressed (silenced) in these five genomes. However, *V. carteri* was exceptional as more than 47% of all complete TRIMs had completely identical LTRs, indicative of recent activity.

To further explore TRIM dynamics, the insertion times of different subfamilies were compared within *G. max*, *P. bretschneideri*, *O. sativa*, *S. moellendorffii* and *V. carteri* (Supplemental Figure S5). Distinct amplification patterns were found between subfamilies in all five genomes. For four genomes (excepting *O. sativa*), one or two subfamilies were dominant, indicating that these TRIMs were likely the most active. For example, in *P. bretschneideri* 2,036 complete PbrRetroS1 (*Pb1*) elements were identified,, 89% of the complete TRIMs. Thus, we postulated that most TRIMs are not extremely active, but that a few subfamilies maintain transpositional activity over long evolutionary timeframes thereby increasing in copy number. To test this hypothesis, we investigated an additional four genomes, *Malus x domestica*, *Z. mays*, *P. trichocarpa* and *P. dactylifera*, as they harbor many complete TRIMs, ranging from 777 to 2,043 (Table 1). We observed similar results to those seen in *P. bretschneideri* (Supplemental Figure S5). Together, these data show differences in transpositional activity between subfamilies and that most TRIMs were likely suppressed/silent or their autonomos retrotransposons were lost or nonfunctional. However, a few subfamilies can remain active over long timeframes resulting in increased copy numbers.

### Putative autonomous retrotransposons and recent transpositions of TRIMs

TRIMs are small elements with no coding capacity, non-autonomous, thus mobilization depends on transposases encoded by other autonomous transposons. However, no autonomous transposon for any TRIM has been reported in plants and the red harvester ant. To identify potential autonomous elements, all 289 TRIM subfamilies were used as queries to search against the 48 plant genomes and GenBank, to find related, but longer elements. For most, 278, no transposase-encoding element was found, but for 11 subfamilies we identified larger, complete elements ranging in size from 3,367 to 8,504 bp, encoding proteins 384-1577 amino acids (aa) in length (Supplemental Table S9). The retroelements can be classified as either Ty1-copia or Ty3-gypsy LTR retrotransposons based on sequence similarity to other retrotransposons. The LTRs of the large retroelements exhibit 79-98% sequence identity with the related TRIMs. Furthermore, the LTR sizes of the TRIMs and their related larger retrotransposons were similar (Supplemental Table S9).

Sequence similarity between the large elements and the TRIMs was not restricted to LTR regions. We identified an 8,504-bp Ty1-copia retrotransposon, OsajLTRA10, in Nipponbare (*Oryza sativa* L. ssp. *japonica*) using 408-bp TRIM OsajRetroS10 as a query. The LTRs of both elements are 115 bp and share 97% sequence identity. OsajRetroS10 also shows 98% and 94% sequence identity with OsajLTRA10 at positions 1-130 and 131-408, respectively, which covers all of OsJRetroS10 (Figure 5A). From this, we infer that OsJRetroS10 is a derivative of OsajRetroA10 via internal deletions with the breakpoint near the 130^th^ nucleotide of OsajLTRA10. There are three complete OsajLTRA10 elements in Nipponbare including OsajLTRA10 on chromosome 1 and other two copies [OsajLTRA10-1 (9,948 bp, on chromosome 9) and OsajLTRA10-2 (5,124 bp, on chromosome 12)]. Sequence alignment of OsajLTRA10 elements and OsajRetroS10 TRIMs reveal that the complete elements contain a 25-bp sequence (CGATCCTA(C/T)AA(G/T)TGGTATCAGAGCC) immediately 5’ of the breakpoint site, and the three OsajLTRA10 elements contained another nearly identical 25-bp sequence immediately 3’ side of the breakpoint site. We refer to this as ‘duplicated internal sequence’ (DIS). The 25-bp DIS were also found in OsaiLTRA10 in 93-11 (*Oryza sativa* L. ssp. *indica*), a close relative of Nipponbare. Additionally, 8-bp (ACCATTGT) and 14-bp (AACAATTGGTATCA) DISs were found in other two TRIM-associated retrotransposons, SmoLTRA4 in *S. moellendorffii* and GmaLTRA2 in *G. max*, however, no DISs were identified in other large retroelements.

**Figure 5.**
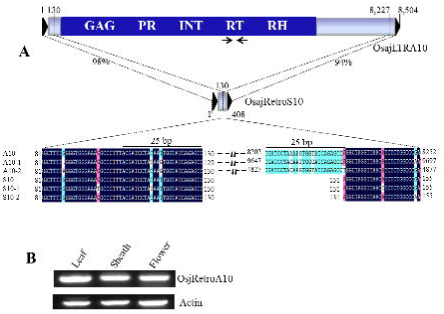
A. OsajRetroS10 and a putative autonomous LTR retrotransposon. OsajRetroS10 is 408 bp and shares high sequence identity with 8,504-bp Ty1-copia retrotransposon OsajLTRA10 in both the LTR and internal regions. OsajLTRA10 contains a duplicated 25-bp sequence, indicated by black lines. **B. RT-PCR analysis of OsajLTRA10**. Primers targeting the conserved domain of the reverse transcriptase (RT) are indicated by arrows.

Among the 11 large LTR retrotransposons, SlyLTRA4, PtrLTRA2, VviLTRA5, PbrLTRA6, CarLTRA1, CarLTRA2 and GmaLTRA2 are likely unable to mobilize TRIMs as their retrotransposon proteins are either short or truncated. Four elements, SitLTRA5, OsajLTRA10, OsiLTRA10 and SmoLTRA4 encode 1409, 1577, 1431 and 1218-aa retrotransposases, respectively, that contain all functional domains for retrotransposition. Thus, these four LTR retrotransposons are putative autonomous elements that can mobilize their related TRIMs. Furthermore, 3, 73, 17 and 13 expressed sequence tags (ESTs) showing sequence similarity with SitLTRA5, OsajLTRA10, OsaiLTRA10 and SmoLTRA4 retrotransposons, respectively, were identified confirming the transcriptional activity of these LTR retrotransposons. As more ESTs were found for OsajLTRA10, we performed RT-PCR analysis to validate the expression of OsajLTRA10 using primers complementary to the reverse transcriptase (RT) domain (Figure 5A). Significant amplification was detected using cDNA from leaf, sheath and flower of Nipponbare and confirms the transcriptional activity of the OsajLTRA10 transposon (Figure 5B).

To gain more insight into the activity of TRIMs, we compared TRIMs from reference genomes for two rice subspecies, *japonica* and *indica*, that diverged ∽0.2-0.4 Mya from either *O. nivara* or *O. rufipogon* (Ma and Bennetzen 2004; Huang et al. 2012), and identified 41 and 31 polymorphic TRIMs in Nipponbare and 93-11, respectively (Supplemental Table S10). All these polymorphic elements were flanked by 5-bp TSDs and absent in the orthologous regions. This suggests that these are newly inserted TRIMs and that transposition of TRIMs may be similar to that of LTR retrotransposons, as both create 5-bp TSDs.

We next conducted PCR to validate the new insertions of OsaRetroS10 for which a putative autonomous retrotransposon was found in both Nipponbare and 93-11 (Figure 5A, Supplemental Table S9). Three pairs of primers targeted to the flanking regions of new insertions sites (Supplemental Figure S6A) were used to amplify DNA from seven rice varieties including four *japonica* (Nipponbare, Kitaaki, Azucena and Moroberkan), three *indica* (93-11, IR36 and IR64) and two AA wild relatives, *O. nivara* and *O. rufipogon*. All three primer pairs yielded expected PCR product sizes in both Nipponbare and 93-11 and the two wild rice species (Supplemental Figure S6B) indicating that these TRIMs were mobilized after the divergence of these two rice subspecies. Interestingly, smaller bands were found in Kitaaki with P1 primers and IR64 with P2 primers. Sequence analysis did not show a deletion in either Kitaaki or IR64, rather an extra complete element and 5-bp sequence were found in the insertion site of Nipponbare and 93-11, respectively. This indicates that OsaRetroS10 may still be active in rice.

## Discussion

### Detection and comparison of TRIMs across the plant kingdom

Due to their diminutive sizes and diverged sequences, TRIMS are difficult to annotate. The first TRIM was identified during analysis of the urease gene using dot plot software (Witte et al. 2001). Since then, other TRIMs have been discovered through comparison of orthologous sequences (Yang et al. 2007) or by PCR cloning experiments (Antonius-Klemola et al. 2006; Kalendar et al. 2008). However, these approaches are time consuming and not suited for genome-wide identification of TRIMs. Computational tools have been developed for *de novo* identification and classification of LTR retrotransposons [e.g. LTR_STRUC (McCarthy and McDonald 2003), LTR-Finder (Xu and Wang 2007), LTRharvest (Ellinghaus et al. 2008) and LTRdigest (Steinbiss et al. 2009)]. However, these tools have limited application for finding TRIMs. For instance, the LTR_STRUC program is inefficient at detecting small retrotransposons (less than 1000 bp), thus, the majority of TRIMs (Supplemental Figure S1A) would be missed. Both LTR-Finder and LTRharvest allow users to define search parameters to find short elements, but will miss diverged elements that lack the primer binding site (PBS) and/or polypurine tract (PPT). LTRdigest requires a retrotransposase sequence, lacking in TRIMs.

In this study, we combined *de novo* annotation and homology-based searches to annotate plant TRIMs. The genomes were initially screened with LTR_Finder to identify TRIMs with typical sequence features. Then, TRIMs identified in other genomes were used to find homologous TRIM sequences missed by LTR-Finder. This combined approach detects more TRIMs than simply using *de novo* annotation and thus provides a good strategy to identify TRIMs in other genomes. For example, of the 11 TRIM subfamilies in *S pimpinellifolium*, only one was detected by LTR_Finder, the other ten were found by homology-searches using TRIMs from *S. lycopersicum* and *C. lanatus*. Furthermore, current annotation tools are not suited for short DNA sequences such as ESTs and genome survey sequences (GSS), whereas homology-based searches can detect TRIMs in these datasets. Our protocol can be used to easily find TRIMs in new sequenced plant genomes. Although TRIM have not been reported in animals, excepting the red harvester ant, this method also works for identifying TRIMs in animals where TRIMs may have been missed by traditional transposon annotations as we found new TRIMs in other ants (Gao et al. unpublished).

Most comparisons of TEs have been limited to closely related species (Ammiraju et al. 2007; Sanyal et al. 2010) or performed at the protein level with conserved transposase domains (Du et al. 2010; Llorens et al. 2011). This is because transposons from distantly related plants are often diverged at the nucleotide level, thus, it is difficult to compare and classify transposons from distantly related genomes, especially for fragmented elements and those that lack transposon proteins.

TRIMs are unusual elements that have been mostly ignored during annotation of plant genomes as only ten TRIM families had been reported in flowering plants thus far. In this study, we used 48 genomes that span ∽610 Myr of plant evolutionary history (Clarke et al. 2011) to identify TRIMs in flowering plants, lycophytes and algae. TRIMs from these species were grouped into 156 TRIM families including 146 novel TRIMs. Of these families, 104 were shared across a range of taxonomic groups. To our knowledge this is the most comprehensive exploration and classification of TRIMs in the plant kingdom. These results provide a valuable resource to the genomics community for identification of homologous TRIM elements in newly sequenced genomes and to gain insight into genome variation mediated by TRIMs. Additionally, the TRIMs can also be developed as molecular markers for detecting genetic diversity (Kwon et al. 2007) and for cloning genes as TRIMs are abundant in many plants and enriched in or near genes (Table 1, Supplemental Table S2).

### Origin and transpositions of TRIMs

No autonomous element has been reported for any of the previously reported TRIMs (Witte et al. 2001; Yang et al. 2007; Kalendar et al. 2008; Gao et al. 2012; Zhou and Cahan 2012). Thus, the evolutionary origin and transposition mechanism of TRIMs remains ambiguous. We found 11 large LTR retrotransposons that share high sequence similarity with specific TRIM LTRs and internal regions and have similarly sized LTRs (Supplemental Table S9, Figure 5). This is the first direct evidence that TRIMs are likely derived from LTR retrotransposons. Duplicated internal sequences (DISs), ranging in size from 8-25 bp in the internal regions of four of the 11 large LTR retrotransposons were observed that may be related to the presumably complex mechanism required to generate TRIMs, although this remains to be confirmed.

To gain insight into the origin of TRIMs, we constructed a phylogenetic tree with the conserved RT domains of the large retrotransposons which revealed that TRIMs are derived from different LTR retrotransposon families and that their origin postdates the emergence of the LTR retrotransposon superfamilies, Ty1-copia and Ty3-gypsy (Supplemental Tables S7 and S9). Notably, the large retroelements identified from the six flowering plants were all Ty1-copia types, whereas, the large retrotransposon from *S. moellendorffii* was a Ty3-gypsy type. It is tempting to speculate that this may reflect an origin for TRIMs from Ty1-copia elements in flowering plants versus Ty3-gypsy elements in *S. moellendorffii*. Additional genome sequencing and annotation is needed to test this hypothesis.

TRIMs do not encode a retrotransposase, therefore their movement depends on proteins produced by other autonomous transposons. Of the 11 TRIM-related LTR retrotransposons (Supplemental Table S9), seven encode short or truncated proteins and are likely nonautonomous LTR retrotransposons. However, four encode full retrotransposases and are putative autonomous elements for TRIMs. Our genome-wide comparisons of TRIMs between two subspecies of *O. sativa* and PCR survey confirmed recent transpositional activity of OsaRetroS10 in *O. sativa*, which contains a related, autonomous LTR retrotransposon.

Of the identified 289 TRIM subfamilies only 11 have related larger LTR retrotransposons. Some full retrotransposons may have been missed due to incomplete genome assemblies. On the other hand, it may reflect selective pressures in plant genomes where transposons are subjected to strong selective pressure to avoid disruption of host genes (Long et al. 2013). However, many TRIMs are highly conserved across species and have likely colonized plants for more than ten million years (Table 1, Figure 1). This leads to questions of how TRIMs can be retained over such long evolutionary times and not removed through mutation or deletion? One strategy may be that TRIMs are small and insert into non-coding regions, such as introns (Supplemental Table S2). Another explanation may be that interactions between TRIMs and their autonomous elements have been lost as the autonomous elements have been removed and thus the TRIMs are no longer mobile. For instance, even though the OsaRetroS10 TRIM is shared between *O. sativa* and a wild rice, *O. brachyantha* (Figure 1), the complete autonomous element was identified only in *O. sativa*, fragments of the autonomous element were found in *O. brachyantha*.

### Unique and evolutionary features of TRIMs

Even though TRIMs are similar in structure to LTR retrotransposons, there are several differentiating features. First and most obvious is their diminutive size. We found that the sizes of more than 77% of the identified TRIMs were less than 1,000 bp, much smaller than the most LTR retrotransposons. Therefore, unlike LTR retrotransposons, the amplification of TRIMs has had less impact on genome expansion. Instead, TRIMs may represent another way to counteract genome expansion accompanying epigenetic silencing (Lisch 2013) and illegitimate recombination (Devos et al. 2002). Notably, the smallest TRIM, CcaRetroS9 in *C. cajan*, was only 233 bp with 52-bp LTRs with 10 complete copies in the genome.

Second, TRIMs are enriched in or near genic regions. We found that an average of 34.1% of the TRIMs in 14 plants were located in or within 1.5 Kb upstream of genes (Supplemental Table S2). We also compared TRIMs from two rice subspecies and found that ∽40% of recently inserted TRIMs were in or near genes (Supplemental Table S11). This suggests that TRIMs more frequently insert, or are retained, into genic regions. Our data also indicates a role for TRIMs in gene evolution based on the following: 1) Recruitment of TRIMs as exons. Some TRIMs were involved in the creation of new exons thereby increasing protein diversity while another TRIMs served as UTRs. UTRs are not translated but they play important roles in stability and localization of the mRNA as well as in regulation of mRNA translation (Mignone et al. 2002). 2) TRIMs contributed to divergence of functional genes. We found 12 and 32 TRIM-related and expressed genes present only in *G. max* and *Z. mays*, respectively, which may represent new genes, or that homolgous genes were either absent or highly diverged in other species. We also found many genes that directly recruited TRIMs as exons (Supplemental Figure S3, 4) which represents a path to the creation of novel genes (Long et al. 2003, 2013). 3) TRIMs were involved in gene acquisition. Many DNA transposons, such as Helitrons, Mutator-like transposable elements (MULEs) and CACTA elements, have been reported to carry fragmented host genes, including both exons and introns (Kapitonov and Jurka 2003; Jiang et al., 2004; Kawasaki and Nitasaka 2004; Pritham and Feschotte 2007). Unlike the DNA-mediated gene capture, TRIMs contain only exonic sequences, similar to a LTR retrotransposon in *Z. mays* and L1 retrotransposon in human. This indicates that the host gene transduction occurred by read-through transcription of retroelements (Jin and Bennetzen 1994; Goodier et al. 2000; Elrouby and Bureau 2010). One gene transduction by a TRIM was reported in *A. thaliana* (Witte et al. 2001), we identified seven TRIM-related transductions in *M. truncatula* and *G. max* (Supplemental Table S6). Our data also provide evidence about retrotranpositions of TRIMs after gene transductions as more than two complete copies were found for five TRIM-containing gene sequences. We found for the first time multiple gene transductions by TRIMs (Figure 2). Taken together with our previous results that TRIMs are likely involved in gene regulation via TRIM-related siRNAs (Gao et al. 2012), TRIMs may represent important contributor in gene evolution and regulation per the early hypothesis that TEs act as “controlling elements” of genes (McClintock 1951).

Third, TRIMs are conserved, retained in plant genomes over long evolutionary timeframes. LTR retrotransposons are dynamic and rapidly diverging sequences (Jiang et al. 2002; Ammiraju et al. 2007; Sanyal et al. 2010), with few exceptions (e.g. centromeric retrotransposons (CRRs) that are shared within grass family (Jiang et al. 2003)). Most plant LTR retroelements are present in only a single genome or in closely related genomes. In contrast, 104 (67%) TRIM families were shared within plant families and/or between distantly related species (Figure 1, Table 1). This may be an underestimate as the other 52 families that are present in only a single plant species may be present in as yet unsequenced species. Additionally, we found numerous TRIMs that inserted more than 5 Mya, whereas, only a handful of LTR retroelements with insertion times older than 5 Myr have been observed. Through comparative analyses, we found 23 and 21 TRIMs located in the orthologous regions of *Z. mays*/*S. bicolor* and *G. max*/*C. cajan*, respectively. Thus, TRIMs are able to colonize and be retained in plants over longer evolutionary period than typical LTR retrotransposons (Vitte et al. 2007). There are a few reasons that could lead to the detection of “old” TRIMs. 1) TRIMs are smaller so there is less opportunity to have nested insertions or truncations resulting in the conservation of ‘old’ elements. 2) Elements in genic or non-genic region evolve differently and as many TRIMs were recruited as exons and they have likely undergone stronger purifying selection (Davidson et al. 2009).

Finally, TRIMs were often organized into tandem arrays. Previous studies have shown that tandem repeats can be functional genomic components. For example, centromeric tandem repeats are essential for formation and maintenance of functional centromeres in both plants and animals (Jiang et al. 2003; Melters et al. 2013). Tandem repeats also found in the human genome and have been linked to many neurodegenerative diseases via gene regulation (Duitama et al. 2014). In this study, we observed tandemly arrayed TRIMs in 35 plant genomes (Supplemental Table S7) indicating that TA-TRIMs are common. LTR retrotransposons are among the most prevalent elements in plants, but tandemly arrayed LTR retrotransposons have not been reported. The mechanism by which TA-TRIMs were created was likely due to homologous recombination between different elements (Devos et al. 2002; Yin et al. 2014). The factors that affect the generations of TA-TRIMs are not clear; however, our data indicate that it is not strongly correlated with TA-TRIM size. Due to the effort required, we manually inspected and validated TA-TRIMs only in *Z. mays* (Supplemental Table S8) where we found evidence for mobilization of TA-TRIMs as seen by multiple copies of TA-TRIMs with different TSDs. As some TA-TRIMs contain more than three internal regions and four LTRs, it was possible that these TRIMs have undergone multiple rounds of sequence exchange and recombination.

In summary, we conducted the most comprehensive analysis of TRIMs thus far and found that TRIMs were distributed and conserved across a range of plant species and were often tandemly arrayed. Our results also suggested that TRIMs appear to be derived from LTR retrotransposons and, in a few species, autonomous LTR retrotransposons were found that likely mobilize TRIMs, although the interactions between TRIMs and the potential autonomous retrotransposons needs to be verified by additional experiments. TRIM are enriched in and near genes and have contributed to genetic novelty, including UTRs, exons and creation of new genes. Thus, from an evolutionary and functional perspective, TRIMs are potentially important sources of genetic novelty that have received scant attention during genome annotation and analysis. Our data provide a holistic view of TRIMs and their unique roles in the plant kingdom and expands our understanding plant genome evolution mediated by LTR retrotransposons.

## Materials and methods

### Plant materials

A total of 10 plant genotypes were used in this study, including an inbred line B73 used for maize genome sequencing project, two wild rice species, *O. nivara* and *O. rufipogon*, and seven cultivated rice, Nipponbare, Kitaaki, Azucena, Moroberkan, 93-11, IR36 and IR64. The seeds of all these plants were planted and grown in the greenhouse at University of Georgia (UGA) with the temperatures set at 30 °C/25 °C (day/night) and a photoperiod at 12 h : 12 h (light/dark). DNAs were extracted from leaves using a CTAB method.

### Plant genome sequences

48 whole genome sequences from a wide evolutionary range of plants were used for annotation of TRIMs, the information for these genomes, gene annotation and availability are shown in Supplemental Table S1. Only the genomes published as of April 1, 2013 were included.

### TRIM annotation

We combined *de novo* annotation and homology-based searches to discover TRIM elements. First, the 48 genomes were analyzed using LTR-Finder (Xu and Wang 2007) with default parameters except that we set a 30 bp and 500 bp of minimum and maximum LTR length, and 30 bp and 2000 bp minimum and maximum distance between 5’ and 3’ LTRs. The output sequences of all “TRIMs” were then manually inspected to discard incorrectly predicted sequences and to determine the exact boundaries of TRIMs. Additionally, all “TRIM sequences” were used as queries to conduct BLASTN and BLASTX searches against GenBank to determine if they contain retrotransposases or other sequences. We used three criteria to define a TRIM element: 1) The element size should be less than 1500 bp; 2) There are at least two complete copies or one complete element and one solo-LTR, and each of the copies should be flanked by different TSDs; and 3) The element should not contain retrotransposase sequences. Second, the TRIMs annotated by LTR-Finder and the previously reported TRIMs in plants (Witte et al. 2001; Yang et al. 2007; Kalendar et al. 2008; Gao et al. 2012) were combined for BLASTN searches against each of the 48 plant genomes and GenBank using different options, including nucleotide collection (nr/nt), reference genomic sequences, expressed sequence tags (ESTs), genomic survey sequences (gss), high throughput genomic sequences (HTGS) and whole-genome shotgun contigs (wgs). The aims of these searchers were to: 1) Identify TRIMs missed by LTR-Finder; 2) Determine if each of the TRIM elements was conserved or species specific. Finally, the TRIMs annotated by LTR-Finder and homology searches were combined to conduct all-against-all BLASTN searches to group these TRIM elements. Because TRIM elements are conserved between related species and homologous elements from same TRIM family may be present in different genomes (Witte et al. 2001; Kalendar et al. 2008; Gao et al. 2012), we used “subfamily” to define TRIMs in each of the plant genomes following a previous publication (Wicker et al. 2007). Furthermore, we grouped all TRIMs into families using the criteria that TRIM elements from different genomes show significant sequence similarity (E value < 1 X 10^-5^) over 5% of the complete element size. These criteria were used to determine if TRIMs were species specific. If no significant hit was found outside the host genome (either the other 47 species or GenBank), the element was considered species specific.

### Estimation of copy number and insertion time of TRIMs

To estimate the copy number and abundance of TRIMs, TRIM elements were used as a custom library to screen the plant genomes with RepeatMasker (http://www.repeatmasker.org) using default parameters with the “nolow” option. We also set a cutoff score greater than 250 and hit sequence size longer than 50 bp.

The insertion times of TRIMs and other LTR retroelements were calculated using the formula: T = K/2r where T represents insertion time, K is average number of substitutions per aligned site and r means an average substitution rate which is 1.3 X 10^^8^^ substitutions per synonymous site per year as suggested by Ma and Bennetzen in grasses (2004) but was also used in soybean and pear (Du et al. 2010; Yin et al. 2014). In this study, we used this substitution rate for other plants because no average substitution rate for transposons is available for many plants including *V. carteri*. 5’ and 3’ terminal repeat sequences of each of LTR retrotransposons were aligned using ClustalW, and the K value was estimated with the Kimura-2 parameter using MEGA 4 software (Tamura et al. 2007).

### Identification of TRIM-related genes and their homologs

To identify TRIM-related genes, a custom perl script was used to screen the RepeatMasker output files from 14 plant genomes against GFF3 annotation files downloaded from Phytozome (http://www.phytozome.net). To avoid duplicated counting, TRIMs that span both exon and intron or upstream and exon were considered a single exon. The proteins of annotated genes from *G. max* and *Z. mays* were extracted and used to search against the proteins in four related genomes, *C. cajan*, *P. vulgaris*, *S. bicolor* and *O. sativa*, from which only genes which proteins show significant sequence similarity (E value < 1X e^-^5^^) with the query proteins were considered homologous genes. If multiply significant hits were detect for some query genes, only the sequence with lowest E value was included.

### PCR and reverse transcription (RT)-PCR analysis

We performed PCR and RT-PCR analysis following previous protocols (Gao et al. 2012). Briefly, the DNAs from cultivated and wild rice and maize were amplified with the corresponding primers (Supplemental Table S12) to validate insertion polymorphisms of a TRIM in rice and TA-TRIMs in maize, respectively. All amplification reactions were done using a MJ Research PTC-200 thermal cycler and the PCR products were purified with QIAquick PCR purification kits (QIAGEN, Venlo, Netherlands) and sequenced by GENEWIZ, Inc. (South Plainfield, NJ). To detect the transcription activity of the rice retrotransposon OsajLTRA10, we collected the leaves and sheath of 4-week old plants and 2-3 cm young spikes from Nipponbare. Total RNAs were isolated using the TRIZOL Reagent (Invitrogen, Carlsbad, CA). Four micrograms total RNA from each sample was converted into single-strand cDNA with reverse transcriptase (Invitrogen, Carlsbad, CA). The cDNA reactions were then diluted 4 to 5-fold, and 2 μL of the diluted cDNA were used as templates for PCR amplifications with the primers targeted to the retrotransposon and actin gene (Supplemental Table S12).

## Acknowledgments

The authors greatly thank Drs. Manyuan Long and Ning Jiang for their critical comments and valuable suggestions. This research was funded by the National Science Foundation grants (IOS0701382 and MCB1026200) to SAJ.

## Disclosure declaration

The authors declare no competing financial interests.

